# Effects of Inosine-5’-Monophosphate Dehydrogenase (IMPDH/GuaB) Inhibitors on *Borrelia burgdorferi* Growth in Standard and Modified Culture Conditions

**DOI:** 10.1101/2024.09.21.614261

**Authors:** Eric L. Siegel, Connor Rich, Sanchana Saravanan, Patrick Pearson, Guang Xu, Stephen M. Rich

**Affiliations:** Laboratory of Medical Zoology, Department of Microbiology, University of Massachusetts, Amherst, MA, 01002, United States of America; (E.L.S.); (C.R.); (S.S.); (P.P.); (G.X.); New England Centre of Excellence in Vector-borne Disease

**Keywords:** Borrelia burgdorferi, Inosine-5’-Monophosphate Dehydrogenase, GuaB, Lyme disease

## Abstract

*Borrelia burgdorferi*’s Inosine-5’-Monophosphate dehydrogenase (IMPDH, GuaB) is a potential therapeutic target. GuaB is not needed for *B. burgdorferi* survival and replication in standard growth media, therefore our ability to test novel drugs against the whole organism is limited to tests using mammalian hosts. Modifications to standard culture conditions that can induce borreliastatic effects of GuaB inhibitors will enhance our ability to evaluate these inhibitors. This study aimed to evaluate basic modifications to standard growth media that can achieve this objective. Effects of GuaB inhibitors (mycophenolic acid, 6-Chloropurine) were compared in standard Barbour-Stoenner-Kelly-II (BSK-II) medium with BSK-II modified to 60% concentration rabbit serum, using *B. burgdorferi* 5A1. Effects were also evaluated with another BSK-II modification with the direct supplementation of hypoxanthine, adenine, and nicotinamide (75 μM each). In standard BSK-II, neither mycophenolic acid nor 6-Chloropurine riboside affected *B. burgdorferi* growth. An ANOVA showed significant, dose-dependent interactions between culture condition and treatment considering modified growth conditions (F = 4.471, P = 0.001). Mycophenolic acid at 250 μM reduced replication by 48 and 50% (P < 0.001 each). 6-Chloropurine riboside was more effective in both conditions at 250 μM, reducing growth by 64 and 65% (P < 0.001 each). Effects of GuaB inhibitors were shown against whole organism *Borrelia burgdorferi* for the first time. Modification of BSK-II medium with physiologically relevant levels of mammalian serum supports replication and induces GuaB inhibitor-mediated suppression of replication. Effects of 6-Chloropurine riboside were demonstrated for the first time, indicating that B. burgdorferi can salvage and phosphorylate these analogue purine derivatives. These molecules can be manipulated to better inhibit replication. Optimisation will allow the assessment of novel Borrelia-specific IMPDH inhibitor molecules in culture for Lyme disease interventions.

**Highlights:**

- Effects of IMPDH inhibitors are shown against whole organism *Borrelia burgdorferi* for the first time.
- Modification of BSK-II medium with physiologically relevant levels of mammalian serum supports replication and induces IMPDH inhibitor-mediated suppression of replication.
- Effects of 6-Chloropurine riboside are demonstrated for the first time, indicating that *B. burgdorferi* can salvage and phosphorylate these analogue purine derivatives. These molecules can be manipulated to better inhibit replication.
- Optimisation of modified BSK-II will allow the assessment of novel *Borrelia*-specific IMPDH inhibitor molecules in culture for Lyme disease interventions.

## 1. Introduction

Lyme disease is the most prevalent vector-borne disease in the United States [1]. Its aetiological agent, *Borrelia burgdorferi*, confers an estimated 476,000 annual cases through *Ixodes* tick bites [2]. Timely antibiotic treatment usually results in complete recovery [3]. However, patients may experience prolonged clinical signs associated with secondary disease of the cardiac, nervous, renal, and other systems [4]. These serious manifestations may worsen when coupled with co-infections of other tick-borne pathogens [5]. New, effective therapeutics targeting human infections and intervention in the *B. burgdorferi* enzootic cycle are needed to address limitations including the possibility of resistance arising to available conventional treatments and incomplete effectiveness.

Nucleotide biosynthesis is a fundamental process to cellular replication, function, and homeostasis [6]. Like many obligate pathogens with compact genomes, *Borrelia* spp. lack the requisite pathways to perform de novo nucleotide biosynthesis [7]. Instead, they must rely on an essential pathway of salvage and interconversion of exogenous precursors to meet replication needs [8]. Limiting steps of similar pathways have been exploited for virulence- and proliferation-disrupting therapeutics for mammalian hosts. One example of successful application was the development of pyrimethamine, which selectively disrupts dihydrofolate reductases of *Plasmodium* spp. and *Toxoplasma gondii* with minimal toxicity to the mammalian host [9,10]. Developing drugs that target nucleotide biosynthesis enzymes requires selective targeting to minimize toxicity and off-target effects. This is because divergence of microbial enzymes from mammalian orthologs complicate binding affinities, thus reducing effectiveness, bioavailability and delivery.

Inosine-5’-Monophosphate Dehydrogenase (IMPDH) is a critical enzyme that catalyzes the limiting, NAD mediated interconversion of inosine-5’-monophosphate (IMP) to xanthosine-5’-monophosphate (XMP). XMP is then converted to guanosine monophosphate (GMP) by GMP synthase for use in DNA/RNA [11]. Mycophenolic acid and ribavirin are examples of IMPDH inhibitors that have proven effective as human cancer, viral, and immune therapies [11].

Differences in catalytic behaviour and active site of mammalian and microbial IMPDHs are well-defined. The binding affinities and effectiveness of available IMPDH inhibitors are largely ineffective or too poorly selective for use in mammalian hosts. For example, mycophenolic acid is shown to inhibit mammalian IMPDHs at much lower concentrations (human 10-20 nM) than that *for B. burgdorferi* ( 7.9 μM) [11]. Therefore, species-specific IMPDHs would be needed to target *B. burgdorferi* without causing serious harm to the host.

*Borrelia* spp. are maintained in distinct environments across its enzootic cycle [12]. These include the tick midgut and the tissues of reservoir and incidental/dead end mammalian hosts. These environments present disparate relative abundances of purines available for scavenge and thus dictate the use of different pathways to utilize these resources in nucleotide synthesis. The *B. burgdorferi* IMPDH (GuaB, encoded by *guaB*) is essential for utilizing scavenged hypoxanthine and adenine [13]. Hypoxanthine is most abundant in blood plasma, available to *B. burgdorferi* while transiently residing during localized infection. Adenine availability may increase with spirochete dissemination to cardiac, dermal, bladder, and other tissues [14]. Mammalian tissues are deficient in guanine, which dictates the essentiality of the *guaB* pathway [13,14]. By comparison, in the tick midgut, only a fitness advantage is seen with *guaB*+ spirochetes relative to knockout strains [13]. The survival of *guaB*-deficient spirochetes in the tick midgut has been associated with the demonstration of direct transport of guanine (deoxy)ribonucleotides for use in (D)RNA synthesis [15]. This would allow *B. burgdorferi* to circumvent its reliance on the *guaB* pathway when enough source of free guanine is present. *guaB*-deficient spirochetes also do not exhibit a defect in vitro (when grown in nutrient-rich Barbour-Stoenner-Kelly-II (BSK-II) medium) [13]. A similar mechanism of guanine utilisation has been hypothesized to explain this.

Solving the crystal structure of *B. burgdorferi’s* GuaB had made it possible to manipulate available IMPDH inhibitors and develop new drugs that may selectively target *B. burgdorferi* [16]. However, these inhibitors cannot be tested in vitro due to the lack of appropriate culture medium. An in vitro assay fills the crucial gap between testing new drugs in silico or against the purified enzyme before moving into the mammalian model. In this report, we describe culture conditions that sustain *B. burgdorferi* growth and induce borreliastatic effects of GuaB-inhibitors. We demonstrate effects using an IMPDH inhibitor that has yet to be assessed against *B. burgdorferi*, 6-Chloropurine Riboside, and mycophenolic acid which has been tested against purified GuaB [17]. This work represents the first instance of assessing the impacts of GuaB inhibitors against *B. burgdorferi* beyond work with the purified enzyme. We considered three growth conditions based off standard BSK-II. We modified this medium with higher concentrations of rabbit serum, with the hypothesis that physiologically relevant conditions can induce effects due to a shift in relative abundance of purine precursors for scavenge. To further gather evidence for this hypothesis, we used a third growth condition, modifying standard BSK-II with the direct supplementation of exogenous hypoxanthine and adenine, in addition to nicotinamide, the precursor to NAD needed for GuaB. With this work, we outline the needs and path for optimising this system to provide the test system for new GuaB inhibitors.

## 2. Materials and Methods

Cultures were prepared from BSK-II medium formulated in-house and supplemented with rifampicin [50 μg/mL), fosfomycin [20 μg/mL], and amphotericin B [2.5 μg/mL] [18]. Cultures of starting concentrations of roughly 300,000 spirochetes/mL were prepared in 1.5 mL volumes using *B. burgdorferi* strain 5A1, a clonal derivative of B31. Three growth conditions were considered: standard BSK-II; BSK-II with added non-hemolysed rabbit serum (Pel-freeze 31125-5) to 60%; and a comparison group of BSK-II with added hypoxanthine (Sigma-Aldrich 68-94-0), adenine (Sigma-Aldrich 73-24-5), and nicotinamide (Sigma-Aldrich 98-92-0) to 75 μM each in Dimethyl sulfoxide (DMSO) as a vehicle (BSK-II/AHN). Technical grade mycophenolic acid (≥ 98%, Sigma-Aldrich 24280-93-1) and 6-Chloropurine riboside (99%, Sigma-Aldrich 5399-87-1) were formulated in DMSO for testing at 125 μM and 250 μM. Controls were vehicle-only (DMSO) groups. The concentration of DMSO was less than 0.5% of culture, which as previously shown to have no harmful effects on growing spirochetes in culture [19]. Solutions were added to culture at day 0 of incubation in a water jacket incubator at 34 °C. Triplicate counts were performed using dark-field microscopy. Preliminary confirmation that BSK-II with higher concentrations of rabbit serum supported growth was performed and is found in the supplemental data.

To compare drug effects between growth conditions, we applied an ANOVA to normalised data, with growth condition and treatment as fixed factors. To obtain a normalisation constant for each growth condition, we calculated the ratio of growth for triplicate counts relative to the mean of respective controls. This accounted for observed differences in absolute growth between groups without deflating observed variance. Pairwise comparisons of treatments between groups were made with Tukey’s HSD correction, focusing on peak exponential growth using R Studio.

## 3. Results

Spirochete replication trends differed slightly between growth conditions. Higher replication rates were generally observed after day 4. Replication rates were greatest in standard BSK-II. No effects of mycophenolic acid or 6-chloropurine riboside at either test concentrations were observed in standard BSK-II (Fig 1A). In contrast, similar dose-dependent effects on replication were observed in BSK-II/60% and BSK-II/AHN with both compounds inducing a flatter exponential pattern at higher concentrations (Fig 1B, Fig 1C). At exponential growth in BSK-II/60% and BSK-II/AHN , *B. burgdorferi* exposed to mycophenolic acid showed a reduction in growth by 48%, 50% (250 μM) and 38%, 9% (125 μM), respectively. 6-chloropurine riboside was ineffective at 125 μM in BSK-II/60% but showed some activity in BSK-II/AHN resulting in a 30% reduction. At 250 μM, 6-Chloropurine riboside was most effective (64%, 65%).

**Figure 1.**
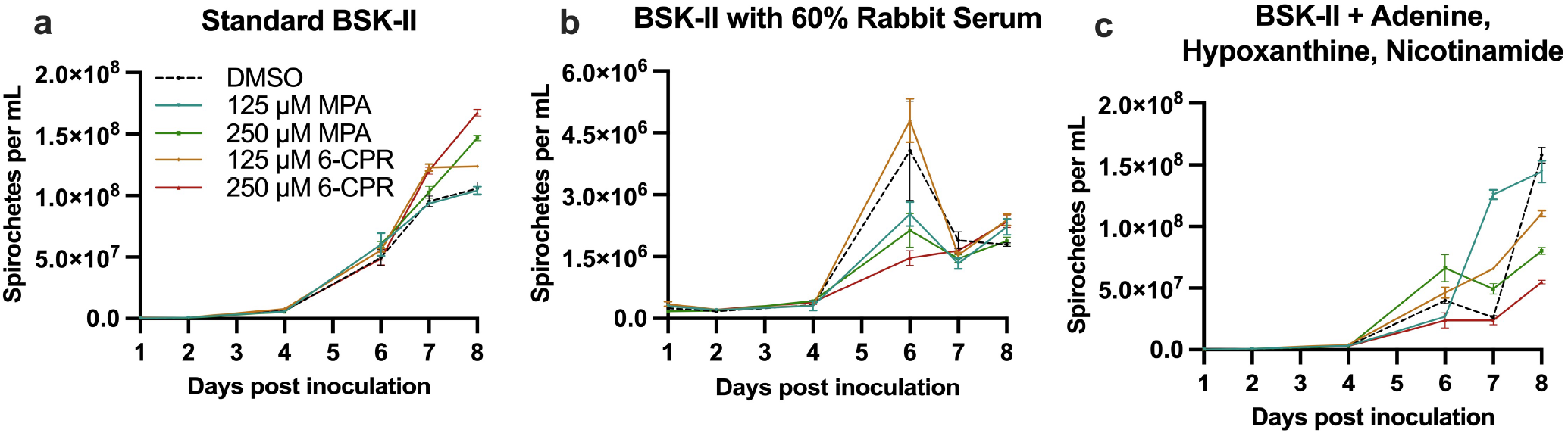
Observed growth of *Borrelia burgdorferi* 5A1 with IMPDH inhibitors under three conditions: (**a**) standard BSK-II; (**b**) modified BSK-II to 60% rabbit serum; (**c**) BSK-II with direct supplementation of 75 μM adenine, hypoxanthine, nicotinamide. Abbreviations: MPA = Mycophenolic acid; 6-CPR = 6-Chloropurine riboside.

A significant interaction between environment and treatment (F = 4.471, P = 0.001) was shown with an ANOVA (Table 1). Effects of 250 μM mycophenolic acid (60% P = 0.0002; AHN P = 0.0006) and 250 μM 6-chloropurine riboside (60% serum P = 0.0007; AHN P = 0.0001) were significantly different from standard BSK-II to BSK-II/60% and BSK-II/AHN (Table 2). 6-chloropurine riboside at 125 μM was ineffective in BSK-II/60% but showed some effects in BSK-II/AHN (P = 0.0243). The effects of mycophenolic acid in BSK-II/60% and BSK-II/AHN at 125 μM were not significant (Figure 2).

**Table 1.** Main effects (growth condition and treatment) and their interaction by ANOVA on cell growth.

| Effect | df | Sum Sq | Mean Sq | F Value | Pr(>F) |
| --- | --- | --- | --- | --- | --- |
| Condition | 2 | 1.186 | 0.5932 | 17.743 | < 0.001 |
| Treatment | 4 | 1.223 | 0.3057 | 9.144 | < 0.001 |
| Condition x Treatment | 8 | 1.196 | 0.1495 | 4.471 | 0.001 |
| Residuals | 30 | 1.003 | 0.0334 |  |  |

**Table 2.** Comparisons for growth condition x treatment interaction with standard BSK-II.

| Treatment | BSK-II/60% Rabbit Serum<br>Estimate (P Sig.) | Direct addition of<br>Hypoxanthine,Adenine,Nicotinamide<br>Estimate (P Sig.) |
| --- | --- | --- |
| 250 µM Mycophenolic Acid | 0.6879 (P = 0.0002) | 0.7072 (P = 0.0001) |
| 125 µM Mycophenolic Acid | 0.3586 (P = 0.0574) | 0.0676 (P = 0.8936) |
| 250 µM 6-chloropurine Riboside | 0.6186 (P = 0.0007) | 0.6325 (P = 0.0006) |
| 125 µM 6-chloropurine Riboside | -0.0650 (P = 0.9010) | 0.4155 (P = 0.0243) |

**Figure 2.**
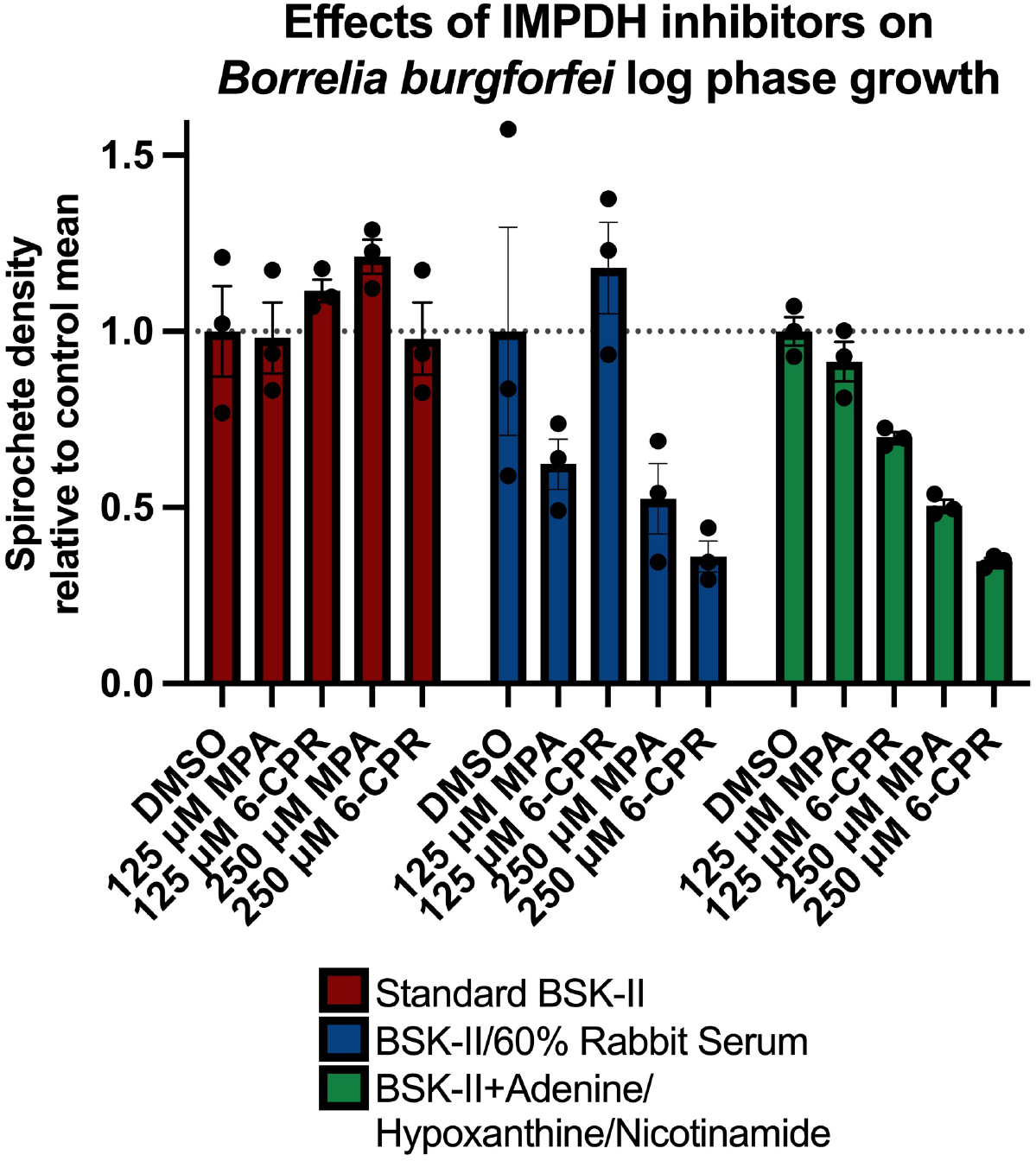
IMPDH inhibitor effects on *Borrelia burgdorferi* growth under three growth conditions at log phase growth. Spirochete concentration is shown relative to the controls of the same growth condition. Significance for the interaction between treatment and growth condition is shown in Table 2. Abbreviations: MPA = Mycophenolic Acid; 6-CPR = 6-Chloropurine Riboside.

## 4. Discussion

This report indicates that IMPDH inhibitors can induce borreliastatic activity in culture, where they should otherwise be ineffective, if the base concentration of BSK-II rabbit serum is increased. The most likely explanation for the induction of drug effects is that higher concentrations of rabbit serum will dilute the BSK-II purine pool for spirochetes to salvage, flooding otherwise accessible free guanine sources with hypoxanthine, adenine, and other unrelated molecules. This would then prompt the requisite increase of *GuaB* expression, which is otherwise likely expressed at low levels in BSK-II, to support replication. In essence, this simulates an exploitative model of the essentiality pattern of *GuaB* observed in the mammalian environment. This hypothesis is supported with the consistency of effects of higher concentrations of mycophenolic acid and 6-Chloropurine riboside were between BSK-II with increased rabbit serum concentration and in BSK-II with direct addition of hypoxanthine, adenine, and nicotinamide. The use of increased serum is likely preferred to direct additions of hypoxanthine, adenine, and nicotinamide as it is more representative of host conditions through spirochete dissemination. The differences observed with growth trends between the two modified conditions warrant further investigation.

We modified BSK-II by increasing the concentration of rabbit serum from standard 6% to 60%. This represents a physiologically relevant concentration to which *B. burgdorferi* is exposed when interacting with the mammalian bloodstream. Plasma (approximately 55% of blood concentration) includes mostly water, proteins, and purines among other molecules accessible for scavenge. *Borrelia* can efficiently salvage purines at low concentrations and would need to be able to use this supply to establish initial presence in the bloodstream. Hypoxanthine is overwhelmingly more abundant in rabbit plasma (5.1 μM, approximately 25x), than adenine and guanine (<0.2 μM) [20]. It may be possible that similar serum concentrations in culture can mimic these conditions and induce g*uaB* expression like what is seen in mammalian infection. While our results suggest that *guaB* expression is upregulated in this condition, it is still unknown how expression is affected by the presence of the other nutrient-rich components in BSK-II. For example, we saw that despite being unable to sustain high levels of replication in the presence of inhibitors, direct addition of adenine and hypoxanthine supported an initial level of growth that was not seen in the 60% serum group. If there are unidentified cues for expression of *guaB* that may govern these differences (present in plasma for example), these could be investigated through means connecting biochemical characterisation with gene expression.

Further investigation into this would require a series of experiments. In the process of doing these, we could further optimize media to provide a system to test new drugs in a translatable environment. The first step would be to titrate BSK-II stocks with rabbit serum at a fine scale. The relative abundance of guanine/hypoxanthine/adenine in each culture needs to be compared to isolated mammalian blood using biochemical means. The hypothesis that components of BSK--II may be a source of excess nucleosides/nucleotides available for scavenge can also be confirmed at this time. Spirochetes can be grown in each condition, and the expression of *guaB* can be compared between conditions and with expression levels in the mammalian host using an already-developed *guaB* reverse transcriptase-quantitative polymerase chain reaction (rt-qPCR) [13]. Once an appropriate medium formulation has been chosen, IMPDH inhibitors can be added to culture and be assessed with inhibition assays. This is important to confirm that the effects we observe are due to GuaB inhibition. Taking 6-chloropurine riboside, for example, this molecule targets the IMP binding pocket of IMPDHs. We cannot rule out that effects could also be compounded by antimetabolite effects at other points in the salvage, transport, and interconversion pathway. This experiment was however an important step as it was unknown whether *B. burgdorferi* would phosphorylate and salvage this analogue-type molecule. With these experiments, we can develop conditions for spirochete growth simulating purine salvage and interconversion in the mammalian host to create an effective test system that could mitigate the need for murine models until optimal compounds are developed.

Other considerations also warrant investigation based on observations in this report. The peak cell concentration in controls grown in BSK-II/60% was not as high as expected based on preliminary experiments (Supplemental data). This could be due to the addition of DMSO, the effects of which have not yet been tested with increased serum concentrations despite having been demonstrated to be acceptable at the concentrations used as a vehicle in standard growth media. Differences in growth could also be attributed to differences between lots of rabbit serum or sources which may vary in exact composition. We also used *B. burgdorferi* 5A1, a laboratory strain that has well-studied properties. It is understood that this strain grows well and is invasive to the range of mammalian hosts of interest. It may be of interest, however, to study effects in naturally occurring strains that are human invasive and perform more intensive biological replicates to accurately characterize effects.

The treatment for Lyme disease is stage dependent. Doxycycline for 10-14 days is the most commonly prescribed antibiotic for Lyme disease without neurologic, cardiac, or joint involvement [3]. Amoxicillin, cefuroxime, and azithromycin may also be used. Treatment is more complex with manifestations associated with secondary disease of major systems [5]. While available antibiotics are usually effective, there remains the need to have more tools available to address clinical disease. It is therefore realistic for GuaB inhibitors to play a major role here. Alternatively, the most effective use case for GuaB inhibitors could be aimed at reducing the prevalence of *B. burgdorferi* in small mammal reservoir hosts, such as the white-footed mouse (*Peromyscus leucopus*) and eastern chipmunk (*Tamias striatus*) which are most abundant in the Northeast, United States [21]. As *GuaB* has been shown to be expressed through several weeks post-infection in murine models, inhibitors in the form of broadcast sprays or feed baits could selectively target *B. burgdorferi* and reduce the spirochete load in the reservoir host. This could result in a reduced prevalence of *B. burgdorferi*-infected hosts on which immature *Ixodes* ticks feed and therefore lower the infection prevalence in ticks for incidental hosts of interest. Doxycycline could theoretically be used for this type of application, however concern about developing resistance in the environment for our most prominently used compound prevents this from being practical. Therefore, GuaB inhibitors are one way that we can meet the need for compounds in this application.

## 5. Conclusions

We demonstrate that BSK-II can be manipulated with higher concentrations of rabbit serum to induce suppression of spirochete growth with GuaB inhibitors, where they would otherwise be ineffective. Effects of GuaB inhibitors are therefore shown in culture for the first time. We also demonstrate for the first time, to our knowledge, effects of 6-chloropurine riboside against *B. burgdorferi*. This work is a foundation for future work that may optimize growth conditions relative to physiological relevance and *guaB* expression patterns. This will allow the assessment of species-specific GuaB inhibitors intended for mammalian administration prior to testing in murine/other animal models. This could provide an easy, accessible alternative to preliminary testing of these molecules.

## Supplementary Materials

S1 Spreadsheet. Growth of *Borrelia burgdorferi* in BSK-II modified with increasing rabbit serum (tab 1); raw counts for GuaB effect calculations (tab 2).

## Author Contributions

Conceptualization, Eric Siegel, Connor Rich, Guang Xu and Stephen Rich; Data curation, Connor Rich and Sanchana Saravanan; Formal analysis, Eric Siegel and Connor Rich; Funding acquisition, Guang Xu and Stephen Rich; Investigation, Eric Siegel; Methodology, Eric Siegel, Connor Rich, Patrick Pearson and Stephen Rich; Project administration, Eric Siegel, Guang Xu and Stephen Rich; Resources, Patrick Pearson and Stephen Rich; Supervision, Connor Rich, Patrick Pearson, Guang Xu and Stephen Rich; Validation, Stephen Rich; Visualization, Eric Siegel; Writing – original draft, Eric Siegel; Writing – review & editing, Eric Siegel, Connor Rich, Sanchana Saravanan, Patrick Pearson, Guang Xu and Stephen Rich. All authors have read and agreed to the final version of the manuscript.

## Institutional Review Board Statement

Not applicable.

## Informed Consent Statement

Not applicable.

## Data Availability Statement

All relevant data are included in the manuscript or supporting files.

## References

1. Kugeler, K.J.; Earley, A.; Mead, P.S.; Hinckley, A.F. Surveillance for Lyme Disease After Implementation of a Revised Case Definition — United States, 2022. MMWR Morb. Mortal. Wkly. Rep. 2024, 73, 118–123, doi:10.15585/mmwr.mm7306a1.

2. Kugeler, K.J.; Schwartz, A.M.; Delorey, M.J.; Mead, P.S.; Hinckley, A.F. Estimating the Frequency of Lyme Disease Diagnoses, United States, 2010–2018. Emerg. Infect. Dis. 2021, 27, 616–619, doi:10.3201/eid2702.202731.

3. Smith, R.P.; Schoen, R.T.; Rahn, D.W.; Sikand, V.K.; Nowakowski, J.; Parenti, D.L.; Holman, M.S.; Persing, D.H.; Steere, A.C. Clinical Characteristics and Treatment Outcome of Early Lyme Disease in Patients with Microbiologically Confirmed Erythema Migrans. Ann Intern Med 2002, 136, 421, doi:10.7326/0003-4819-136-6-200203190-00005.

4. Cardenas-de La Garza, J.A.; De La Cruz-Valadez, E.; Ocampo-Candiani, J.; Welsh, O. Clinical Spectrum of Lyme Disease. Eur J Clin Microbiol Infect Dis 2019, 38, 201–208, doi:10.1007/s10096-018-3417-1.

5. Lantos, P.M.; Wormser, G.P. Chronic Coinfections in Patients Diagnosed with Chronic Lyme Disease: A Systematic Review. The American Journal of Medicine 2014, 127, 1105–1110, doi:10.1016/j.amjmed.2014.05.036.

6. Lane, A.N.; Fan, T.W.-M. Regulation of Mammalian Nucleotide Metabolism and Biosynthesis. Nucleic Acids Research 2015, 43, 2466–2485, doi:10.1093/nar/gkv047.

7. Fraser, C.M.; Casjens, S.; Huang, W.M.; Sutton, G.G.; Clayton, R.; Lathigra, R.; White, O.; Ketchum, K.A.; Dodson, R.; Hickey, E.K.; et al. Genomic Sequence of a Lyme Disease Spirochaete, Borrelia Burgdorferi. Nature 1997, 390, 580–586, doi:10.1038/37551.

8. Pettersson, J.; Schrumpf, M.E.; Raffel, S.J.; Porcella, S.F.; Guyard, C.; Lawrence, K.; Gherardini, F.C.; Schwan, T.G. Purine Salvage Pathways among Borrelia Species. Infect Immun 2007, 75, 3877–3884, doi:10.1128/IAI.00199-07.

9. Lapinskas, P.J.; Ben-Harari, R.R. Perspective on Current and Emerging Drugs in the Treatment of Acute and Chronic Toxoplasmosis. Postgraduate Medicine 2019, 131, 589–596, doi:10.1080/00325481.2019.1655258.

10. Ferone, R.; Burchall, J.J.; Hitchings, G.H. Plasmodium Berghei Dihydrofolate Reductase. Isolation, Properties, and Inhibition by Antifolates. Mol Pharmacol 1969, 5, 49–59.

11. Hedstrom, L. IMP Dehydrogenase: Structure, Mechanism, and Inhibition. Chem. Rev. 2009, 109, 2903–2928, doi:10.1021/cr900021w.

12. Helble, J.D.; McCarthy, J.E.; Hu, L.T. Interactions between Borrelia Burgdorferi and Its Hosts across the Enzootic Cycle. Parasite Immunology 2021, 43, e12816, doi:10.1111/pim.12816.

13. Jewett, M.W.; Lawrence, K.A.; Bestor, A.; Byram, R.; Gherardini, F.; Rosa, P.A. GuaA and GuaB Are Essential for B Orrelia Burgdorferi Survival in the Tick-Mouse Infection Cycle. J Bacteriol 2009, 191, 6231–6241, doi:10.1128/JB.00450-09.

14. Spector, R. Hypoxanthine Transport through the Blood-Brain Barrier. Neurochem Res 1987, 12, 791–796, doi:10.1007/BF00971517.

15. Lawrence, K.A.; Jewett, M.W.; Rosa, P.A.; Gherardini, F.C. Borrelia Burgdorferi Bb0426 Encodes a 2′-deoxyribosyltransferase That Plays a Central Role in Purine Salvage. Molecular Microbiology 2009, 72, 1517–1529, doi:10.1111/j.1365-2958.2009.06740.x.

16. McMillan, F.M.; Cahoon, M.; White, A.; Hedstrom, L.; Petsko, G.A.; Ringe, D. Crystal Structure at 2.4 Å Resolution of Borrelia Burgdorferi Inosine 5’-Monophosphate Dehydrogenase: Evidence of a Substrate-Induced Hinged-Lid Motion by Loop 6,. Biochemistry 2000, 39, 4533– 4542, doi:10.1021/bi992645l.

17. Zhou, X.; Cahoon, M.; Rosa, P.; Hedstrom, L. Expression, Purification, and Characterization of Inosine 5′-Monophosphate Dehydrogenase from Borrelia Burgdorferi. Journal of Biological Chemistry 1997, 272, 21977–21981, doi:10.1074/jbc.272.35.21977.

18. Barbour, A.G. Isolation and Cultivation of Lyme Disease Spirochetes. Yale J Biol Med 1984, 57, 521–525.

19. Lynch, A.; Pearson, P.; Savinov, S.N.; Åli, A.Y.; Rich, S.M. Lactate Dehydrogenase Inhibitors Suppress Borrelia Burgdorferi Growth In Vitro. Pathogens 2023, 12, 962, doi:10.3390/pathogens12070962.

20. Eells, J.T.; Spector, R. Determination of Ribonucleosides, Deoxyribonucleosides, and Purine and Pyrimidine Bases in Adult Rabbit Cerebrospinal Fluid and Plasma. Neurochem Res 1983, 8, 1307–1320, doi:10.1007/BF00964000.

21. Williams, S.C.; Little, E.A.H.; Stafford, K.C.; Molaei, G.; Linske, M.A. Integrated Control of Juvenile Ixodes Scapularis Parasitizing Peromyscus Leucopus in Residential Settings in Connecticut, United States. Ticks and Tick-borne Diseases 2018, 9, 1310–1316, doi:10.1016/j.ttbdis.2018.05.014.

